# CReSCENT: CanceR Single Cell ExpressioN Toolkit

**DOI:** 10.1101/2020.03.27.012740

**Authors:** Suluxan Mohanraj, J. Javier Díaz-Mejía, Martin D. Pham, Hillary Elrick, Mia Husić, Shaikh Rashid, Ping Luo, Prabnur Bal, Kevin Lu, Samarth Patel, Alaina Mahalanabis, Alaine Naidas, Erik Christensen, Danielle Croucher, Laura M. Richards, Parisa Shooshtari, Michael Brudno, Arun K. Ramani, Trevor J. Pugh

**Author notes:** The authors wish it to be known that the first two authors should be regarded as joint First Authors.

## Abstract

**CReSCENT:** CanceR Single Cell ExpressioN Toolkit (https://crescent.cloud), is an intuitive and scalable web portal incorporating a containerized pipeline execution engine for standardized analysis of single-cell RNA sequencing (scRNA-seq) data. While scRNA-seq data for tumour specimens are readily generated, subsequent analysis requires high-performance computing infrastructure and user expertise to build analysis pipelines and tailor interpretation for cancer biology. CReSCENT uses public data sets and preconfigured pipelines that are accessible to computational biology non-experts and are user-editable to allow optimization, comparison, and reanalysis for specific experiments. Users can also upload their own scRNA-seq data for analysis and results can be kept private or shared with other users.

## INTRODUCTION

Recent advances in single-cell RNA-sequencing (scRNA-seq) have enabled the measurement of expression levels of thousands of genes across thousands of individual cells (1). scRNA-seq technologies can be used to identify cell subpopulations with characteristic gene expression profiles in complex cell mixtures, including both cancer and non-malignant cell types within tumours. A typical computational pipeline to process scRNA-seq data involves mapping of reads against a transcriptome, quality control (QC), normalization, dimension reduction, cell clustering, cell cluster labeling, and detection of differentially expressed genes (DEGs). Diverse visualization tools can be used to explore results from these steps, such as t-distributed stochastic neighbor embedding (t-SNE) or Uniform Manifold Approximation and Projection (UMAP) plots for dimension reduction and cell clustering. QC and differential gene expression can be visualized using violin plots. Opportunities to improve scRNA-seq data analysis include making user-friendly and standardized end-to-end analysis pipelines readily available and the ability to tailor analysis parameters to account for atypical gene expression in cancer cells due to underlying genome alterations.

In the last five years, the number of algorithms and packages available for the various components of scRNA-seq analysis has increased considerably (2, 3). In Bioconductor alone, the number of packages for single-cell analysis has increased from two in 2016 to 50 in 2019 (2). This, while providing many options, leaves users with the difficult task of deciding which tools and parameters they should use. Benchmarking studies (4–6) help to facilitate those decisions, however, some analyses require high-performance computing infrastructure and intermediate to advanced levels of computational expertise, limiting their use. Since scRNA-seq technologies are being widely used across various fields of research, standardized analysis pipelines are needed to process data for comparison and integration of data sets. This is particularly relevant for comparison and integration of data across cancer research studies, where scRNA-seq data are now routinely generated from a diversity of patients, cancer types, tissue types, treatment protocols, and cell selection methods.

In this study, we describe CReSCENT: CanceR Single Cell ExpressioN Toolkit (https://crescent.cloud), an intuitive and scalable web portal for standardized analysis and exploration of scRNA-seq data from cancer studies. CReSCENT is accessible through a web-browser and provides these tools without the need for extensive bioinformatics expertise or access to high-performance computing infrastructure. CReSCENT pipelines are end-to-end, removing the need for users to spend time learning how to integrate various independent tools. CReSCENT is populated with multiple public data sets and preconfigured pipelines that are user-configurable, enabling optimization, comparison, and reanalysis of scRNA-seq data. Users can also upload their own experimental scRNA-seq data for analysis and can opt to keep it private, share it with specific users, or make it public. CReSCENT’s interactive data visualizations allow users to deeply explore features of interest without re-running pipelines, as well as overlay additional, custom meta-data such as cell types or T cell receptor sequences. In addition, vector format plots and tables are downloadable for offline analysis and publication purposes.

## METHODS

### Data

For scRNA-seq analysis, CReSCENT requires a gene expression matrix in Matrix Market format (MTX) as input. Compared to gene-by-barcode text files, the MTX format requires less storage space for sparse matrices where many elements are zeros, as is often the case for scRNA-seq data sets. CReSCENT provides one-line-command scripts to convert gene-by-barcode text files to MTX. Metadata for individual cells can also be uploaded to the portal to visualize and colour cells, for example, using results from orthogonal assays, sample annotations, cell annotations, including cell types, and T and B cell receptor sequences from single cells. The metadata format is a tab-separated file with cell barcodes in the first column and appropriate values in the subsequent columns. Two types of variables can be contained in the metadata: ‘group’ for categorical variables, like cell types, and ‘numeric’ for continuous values, like orthogonal assay measurements.

### Analysis

CReSCENT uses Seurat (7, 8), a single-cell analysis R toolkit, to define and configure a standardized scRNA-seq pipeline. The steps included in the pipeline that CReSCENT currently provides are QC, normalization, dimension reduction, cell clustering, cell cluster labelling, and differential gene expression detection. QC filters remove low-quality cell data from the analysis, such as data from dying cells (9). Normalization is necessary to remove technical biases, and CReSCENT allows the user to choose between three normalization options: i) a traditional Log-scale normalization followed by a search of variable genes and data scaling, ii) SCTransform, which provides a framework for the normalization and variance stabilization of molecular count data and reportedly improves common downstream analytical tasks including variable gene selection (7), or iii) the use of pre-normalized measurements (e.g. transcripts per million). Dimension reduction (principal component analysis) identifies the dimensions with the highest variation and uses them for cell clustering.

For cell clustering, CReSCENT provides Seurat’s clustering algorithm, which outperforms other methods in benchmarking studies (4, 6). Non-linear dimension reduction techniques (t-SNE and UMAP) are used to visualize cells in a two-dimensional space and colour them according to features of interest, such as cell clusters, expression of genes of interest, or cell metadata. Finally, DEGs per cluster are detected by comparing the expression of each gene in a cluster to the expression of the same gene in the rest of the cell clusters. For this step we used Seurat’s function FindAllMarkers() with most parameters set to defaults. We specified that only genes detected in a minimum of 25% of cells (parameter ‘min.pct=0.25’) enter the DGE to speed up the function by not testing genes that are very infrequently expressed. Another parameter that we adjusted is the pseudocount to add to averaged expression values when calculating logFC (Seurat’s default=1); instead we use 1 / number of cells in the dataset (10). The top DEGs are shown in t-SNE, UMAP and violin plots and a full list of DEGs is provided as downloadable tables.

### Output

CReSCENT’s graphical user interface (GUI) (Figure 1A) allows users to interactively colour cells on-the-fly using specific features including candidate genes of interest or custom metadata without having to rerun the analysis or reload the R object. CReSCENT provides four types of interactive visualizations: QC violin plots, dimension reduction plots (t-SNE and UMAP), and gene expression violin plots. Within the QC menu, violin plots show the distributions of the configurable cell QC metrics before and after the filtering of cells by those metrics (Figure 1B). This includes the number of genes, the number of reads, the percentage of reads mapped to mitochondrial genes, and the percentage of reads mapped to ribosomal protein genes. These QC metrics can also be viewed individually as gradients in UMAP plots (Figure 1C). All plots in CReSCENT are interactive and the user can zoom into groups of cells or move the mouse cursor over points representing cells to obtain information such as the cell barcode or value of the attribute in the plot. By default, the dimension reduction plots show cells coloured according to cell clusters defined by gene expression using CReSCENT’s pipeline (Figure 2A). Alternatively, these plots can be coloured using the expression of a gene of interest using the ‘Search’ function or using the top DEGs identified by CReSCENT’s pipeline (Figure 2B). The DEG measurements across clusters can be also visualized using violin plots (Figure 2C). Additionally, dimension reduction plots can be coloured using cell metadata provided by the user. Two types of metadata can be used: i) discrete (‘group’) variables such as cell type annotations, which are coloured using solid tones (Figure 2D) or ii) continuous (‘numeric’) variables, such as gene expression measurements from orthogonal assays, which are represented using colour gradients (Figure 2E). Interactive plots, vectorized figures, text tables, and R objects are all downloadable through the GUI for further offline analysis.

**Figure 1.**
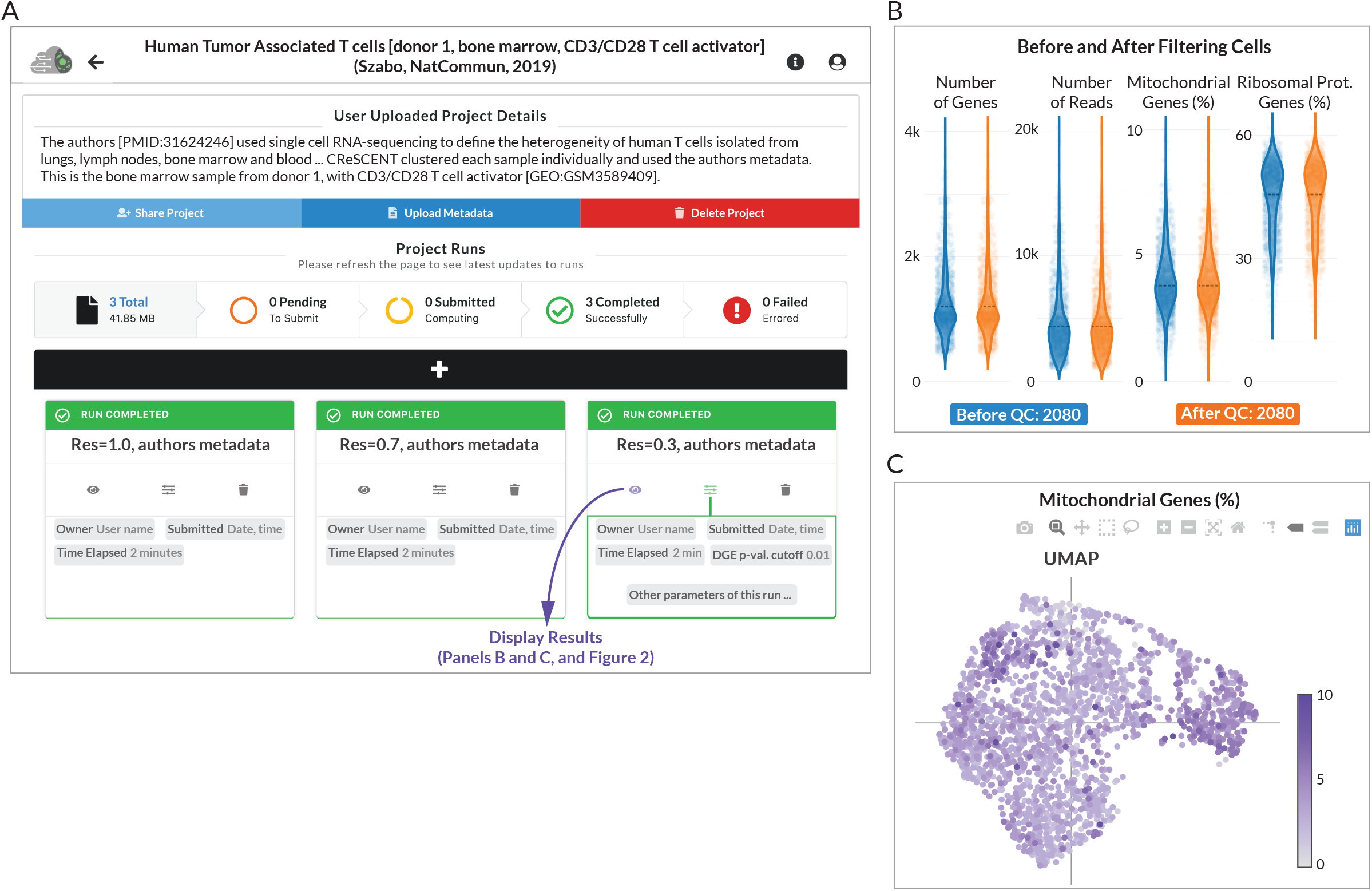
CReSCENT’s web portal run control menu. A) Control menu of a CReSCENT project called Human Tumour-Associated T cells. Users can see details of the project and controls to share the project with other CReSCENT users, upload cell metadata, delete the project, and view number and status of runs in the project. Runs within a project use the same input dataset (i.e. the same scRNA-seq measurements) but parameters of the run change between runs. For example, this panel shows three runs using different clustering resolutions:1.0 (left), 0.7 (middle) and 0.3 (right). Each run box has three controllers: visualization of results, display of the run parameters and deletion of the run. B) QC results from the run with resolution of 0.3. Each dot represents a cell and the violins represent the distribution of one out of four QC metrics. Distributions are shown for cell populations before and after filtering cells by each QC metric. In this run, no cells were filtered out. C) UMAP plot showing one of the four QC metrics (percentage of mitochondrial genes). The other three metrics can be visualized by selecting them from a drop-down menu in the results control panel (shown in Figure 2). Interactive visualization tools allow the user to zoom-in/out and select data from the plots.

**Figure 2.**
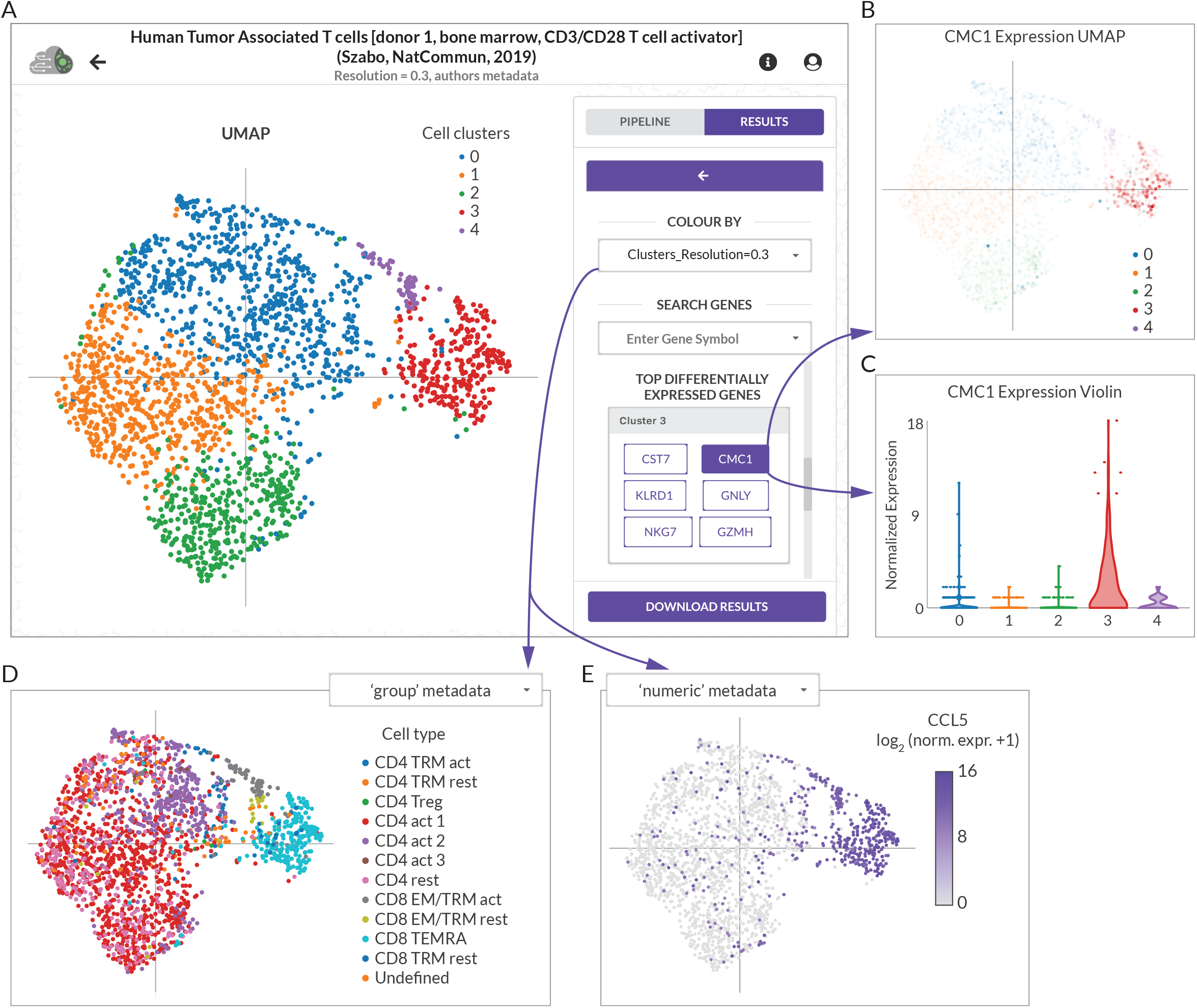
CReSCENT’s web portal visualizations menu. A) Control menu of a CReSCENT project called Human Tumour-Associated T cells, showing a UMAP with the five cell clusters detected by the CReSCENT pipeline. The right-hand menu allows the user to switch between plot types (UMAP, t-SNE or violin) and between the type of data to be plotted (e.g. cell clusters, DEG’s expression or metadata). B) UMAP of CMC1, which is a DEG from cluster 3. Each dot denotes an individual cell and the opacity of the dot corresponds to the expression of CMC1 in that cell. C) Violin plot representation of CMC1 expression across the five cell clusters detected by CReSCENT shows higher expression of the gene in cluster 3. D) UMAP coordinates obtained by CReSCENT’s pipeline are used to show cell metadata provided by the user for cell types (‘group’ discrete categories). E) Similar to D but showing other types of metadata provided by the user for expression of gene CCL5 measured by an orthogonal method with ‘numeric’ continuous values.

### Computing time benchmark

CReSCENT uses Seurat’s parallelization via the R library, ‘future’. We used a scRNA-seq dataset with 68,579 human peripheral blood mononuclear cells (PBMCs) (10) to determine CReSCENT’s pipeline running times. First, we randomly subsampled 60,000 cells from the 68,579 cells dataset, then randomly removed 10,000 cells five times in a stepwise manner to obtain between 50,000 and 10,000 cells, then removed 5,000 cells and finally 4,000 cells to finish with 1,000 cells. For each subsample, we ran the CReSCENT pipeline with default parameters using each of the following number of cores for ‘future’: 1, 2, 4, 10, 20, 30, 40, 50, 60, 70, 75. The procedure, including subsampling and running the CReSCENT pipeline, was repeated three independent times and an average and standard deviation of computing times was calculated for each combination of number of cells and number of cores. All computing times were calculated using an Intel Xeon Processor (Skylake) 2GHz Virtual Machine with 75 cores and 182.6GB RAM.

## IMPLEMENTATION

### Applications

CReSCENT scRNA-seq applications are made in R v3.6.1 (11) using Seurat v3.1.1 (8), ggplot2 v3.2.1 (12), and in-house functions. These workflows are containerized and described below and can also be used as stand-alone one-line-command scripts in R. For reproducibility purposes, CReSCENT generates downloadable files ordered into folders. One of those folders, called LOG_FILES, will contain the following files that the user can use for reproducibility: i) *UsedOptions.txt file with the one-line-commands used for the run, and ii) *RSessionInfo.txt with the R and R library versions used in the run. A copy of the R one-line-command script used in the run is included in the root of the Download Results file provided by our web application. All our software is provided under the GPL-3.0 licence.

### Software Architecture

CReSCENT is designed to enable ease-of-use with the entire portal represented by three pages: ‘Projects’, ‘Project Runs’, and ‘Visualizations’. CReSCENT’s software architecture consolidates a variety of web technologies to achieve this user-friendly interface for scRNA-seq analyses. In Figure 3, we outline these web technologies and their interactions within the CReSCENT architecture. The front end is built using the JavaScript frameworks React (13) and Redux (14) with Nginx (15) serving them in production (Figure 3A and 3B). File storage is supported by MinIO (16) and project and run documents are stored in MongoDB (17) (Figure 3C). The data integration layer, GraphQL (18), facilitates the interactions between these technologies. In the back end, Express (19) is responsible for submitting containerized workflows and spawning Python (20) processes that rapidly query gene expression across large gene-by-cell matrices (Figure 3D). The entire web application is containerized with Docker (21) for accessible deployment.

**Figure 3.**
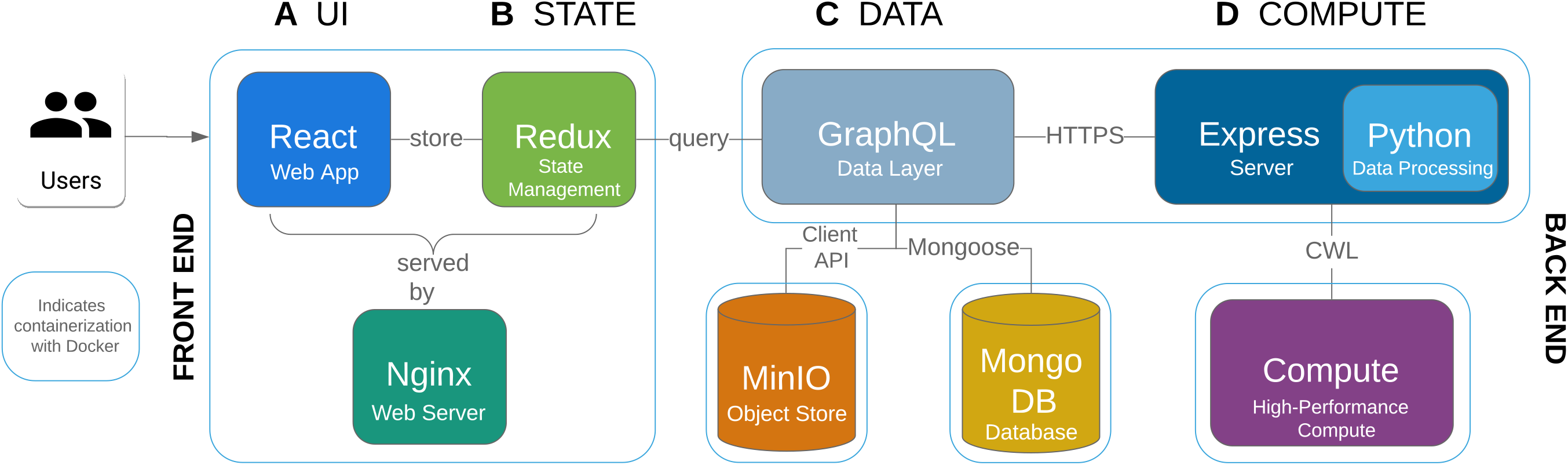
CReSCENT’s web portal technologies and their interactions. Overview of the technology components utilized in developing CReSCENT. Blue lines indicate Docker containers. Files are uploaded through the React front end (A and B) and saved into MinIO and MongoDB (C), parameters are sent to the Express server, and the CWL job is sent to compute (D). Once results are available, project and run documents are updated in MongoDB, results are visualized in the front end with gene expression queries processed by a Python back end, and results are served as a zipped download through MinIO. Interactions are facilitated by GraphQL.

### Data Layer

GraphQL consolidates MinIO and MongoDB technologies into a graph schema and serves an application programming interface (API) to the user interface. The modular graph structure currently includes users, projects, and runs. Only necessary data is queried from the graph, resulting in a more performant approach than commonly used REST APIs. For instance, the structure of a GraphQL query is resolved server-side, resulting in a single client call and server response. This is in contrast to REST APIs, where multiple calls would be made to the server to resolve the resulting data for the client. Using a graph definition also allows for simple versioning and flexible deployment. Furthermore, GraphQL allows for the integration of multiple scRNA-seq data sources as well as project organization via cancer type ontologies, making it an appropriate tool for CReSCENT’s future applications.

### Containerized Compute

The Common Workflow Language (CWL) (22) is a standardized pipeline language used to describe containerized workflows in a portable, reproducible, and scalable fashion. CReSCENT uses CWL to provide users pipeline customizability and to organize jobs within the back end. CWL pipelines are written by mapping inputs and outputs of a workflow. CWL is powerful within scRNA-seq analyses due to the increasing number of dependencies required to install and run these analyses. CReSCENT submits customized jobs through the Express server to run CWL workflows in a high-performance computing environment within Docker or Singularity (23) containers.

### Interactive Visualizations

The JavaScript graphing library Plotly (24) is used to create CReSCENT’s interactive visualizations. A back end written in Python provides QC data and gene expression levels. Barcoded cells are grouped by Seurat clusters or metadata attributes for display. To responsively return normalized gene expression data from up to 100,000 cells, information is stored in Loom files built on the HDF5 standard for storing sparse datasets, and extracted with the Python package LoomPy (25).

## RESULTS

### Human Tumour-Associated T cells

To show how CReSCENT may be used for the analysis of scRNA-seq data, we performed an analysis of a publicly available human tumour-associated T cell scRNA-seq data set, containing data from the blood, lymph nodes, lungs and bone marrow from two donors, with or without CD3/CD28 T Cell Activator stimulation (26). Using one of the samples from this dataset, from the bone marrow of one donor, with CD3/CD28 T cell activation, we performed multiple runs of the CReSCENT pipeline, each with a different cell clustering resolution (1.0 [default], 0.7 and 0.3). Each run and its parameters can be viewed independently in the Project Runs page (Figure 1A), where the status of the run (e.g. run completed) is indicated above the run name. The total number of runs for a project, as well as the number of pending, submitted, completed, and failed runs are also indicated here. From the Project Runs page, additional metadata can be uploaded, and the project can be shared with other CReSCENT users or deleted.

We show QC violin plots with the distribution of the number of genes, the number of reads, percentage of mitochondrial genes, and percentage of ribosomal protein genes measured in the sample before (blue) and after (orange) QC; the average values for each of these metrics is indicated by a dotted black line (Figure 1B). The initial analysis included 2080 cells, all of which were retained after QC (Figure 1B). The same QC metrics can be visualized using UMAP plots. For example, in Figure 1C each dot represents a cell and the percentage of mitochondrial genes measured in that cell correlates to the purple colour intensity of the dot. In Figure 2, we show the results of our analysis using a cell clustering resolution = 0.3, differential gene expression and metadata visualization. At this resolution, we found five cell clusters and CReSCENT’s GUI displays the top DEGs for each cluster on the right-hand side menu (Figure 2A). For example, cluster 3 showed significantly higher expression of the CMC1 gene compared vs. the rest of the cells in other clusters (corrected p-value = 1.189e-122, average fold change = 3.25). Thus, we listed CMC1 as a DEG for cluster 3. This is reflected by both the intensity of the cluster’s red colour in Figure 2B and the dominant distribution of the red violin in Figure 2C when CMC1 is selected from the DEGs menu. Other DEGs within cluster 3 include CST7, NKG7, KLDR1, GNLY, and GZMH, several of which have been previously associated with cytotoxic T cell function (27). This coincides with the analysis from the original authors of this dataset (26), who identified cells belonging to cluster 3 enriched for CD8+ effector memory CD45RA (CD8_TEMRA_) cells (Figure 2D) and expression of CCL5 (Figure 2E), which is a marker of CD8+ TEM cells (26, 28). Together, this demonstrates the ability of the CReSCENT framework to recapitulate findings from original publications and support reproducible single-cell science.

### Computing time benchmark

Using subsamples of a scRNA-seq dataset of 68,579 PBMCs, we found that, in general, using multicore processing reduced computing times between 2 and 4 times compared against using a single core for the same task (Figure 4). We found that using 4 to 10 cores reduced computing times compared with fewer cores, but we did not observe further improvements beyond 10 cores (Figure 4). Interestingly, we found a slight increase in the computing time as the number of cores increases. In practice the sizes of computing problems scale with the amount of available resources. However, if a problem only requires a small amount of resources, it is not beneficial to use a large amount of resources to carry out the computation. This is because as the number of processors increases, computational work per processor decreases, but communication time typically increases, and generally degrades parallel performance (29). Thus a more reasonable choice is to use small amounts of resources for small problems and larger quantities of resources for big problems. Based on our results, CReSCENT’s pipeline automatically allocates 4 cores for runs with <30,000 cells and 10 cores for runs with >= 30,000 cells, to avoid ‘saturation’ of available CPU cores. CReSCENT’s current implementation processes 1,000 cells in ~1 minute, 30,000 cells in ~19 minutes, and 68,000 cells in ~45 minutes (Figure 4). Using the same PBMC subsamples, we measured the time to query and visualize the expression of a single gene in CReSCENT’s GUI. CReSCENT was able to perform gene query and visualization in ~3 seconds for 1,000 cells, ~8 seconds for 30,000 cells, and ~13 seconds for the entire 68,579 cells dataset.

**Figure 4.**
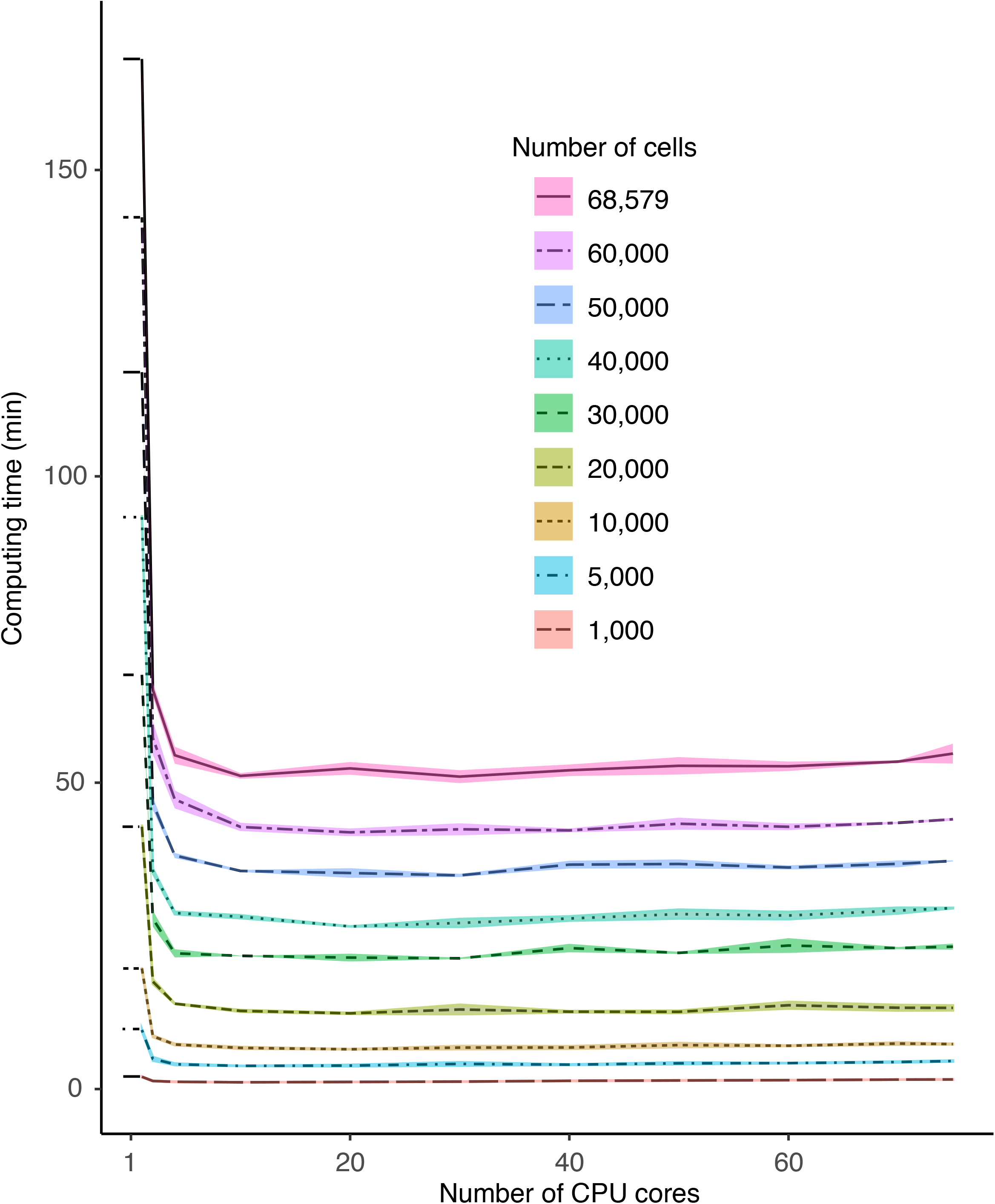
Computing time benchmark. Computing times of runs of the CReSCENT pipeline using either a scRNA-seq dataset of 68,579 PBMCs or random stepwise subsamples of 1,000 to 60,000 cells from the full dataset. The full dataset or each subsample were run three independent times through the pipeline. The number of cores used for each run is indicated in the x-axis and the run time is plotted in the y-axis. Each line represents the average computing time used to process either the full dataset or each of the subsamples in three replications. Each line ribbon represents one standard deviation from the same three replications analyses.

## DISCUSSION

CReSCENT provides a standardized, end-to-end scRNA-seq analysis pipeline that is accessible to users of all bioinformatic expertise levels. Computing time benchmarks showed that CReSCENT is capable of performing cell clustering and DEG detection from raw read counts of 1,000 cells in approximately 1 minute and 68,000 cells in 45 minutes. Importantly, the application is easily accessed via a web browser through a simple and intuitive interface. Software dependencies within CRESCENT are abstracted through containerized workflows resulting in reproducible and scalable analyses. This eliminates the need for users to spend time learning bioinformatic analytical techniques or how to consolidate various analytical tools to achieve their scRNA-seq research goals. Furthermore, CReSCENT provides interactive visualization tools allowing exploration and querying of large data sets in real time. Responsive querying is achieved through Python and Loom file format implementations which are highly scalable. This allows users to find genes of interest pertaining to their experiment as well as explore data-driven, differentially expressed genes in a matter of seconds. This is in contrast to native R sessions where analyses are stored in R objects that can take several minutes to reload into memory, depending on the size of the dataset. Typical scRNA-seq analyses are also rerun frequently to query genes of interest, add new cell metadata, and tune parameters. CReSCENT is designed to streamline and automate these tasks.

We drew inspiration from other single-cell visualization platforms, such as the Broad Institute’s Single Cell Portal (30), and added standardized analysis capabilities. We continue to curate and implement applications and software based on benchmark studies (4–6). In addition to the development of visualization and analytic pipelines, CReSCENT is an ongoing project to collect, store and make available high-quality, cancer-specific scRNA-seq datasets in a standardized format, as well as valuable reference sets from healthy tissues. Currently, in the system, we have deployed pre-processed scRNA-seq data from ~150,000 cells in 12 cancer and one healthy reference tissue datasets.

At this stage, CReSCENT provides several cancer datasets, however, our pipelines continue to be standard for any type of tissue. In the future, CReSCENT will expand to include additional collections of cancer and reference scRNA-seq datasets, and the implementation of the following applications: integration of multiple scRNA-seq data sets, cell type labeling, detection of copy number variations, and alternative cell clustering methods. We expect that these cancer-specific applications will contribute to determining more accurate empirical parameters for cancer studies. We will continue to develop CReSCENT as a versatile scRNA-seq analysis and visualization platform to help cancer researchers accelerate accessible and reproducible single-cell science.

## AVAILABILITY

This web application is free and open to all users and there is no login requirement. The CReSCENT GUI and cancer scRNA-seq data sets are accessible online at https://crescent.cloud with user-friendly documentation available at https://pughlab.github.io/crescent-frontend/. CReSCENT is an open-source initiative with one-line-command scripts and containerized workflows available at https://github.com/pughlab/crescent and the portal code repository available at https://github.com/pughlab/crescent-frontend.

## ACKNOWLEDGEMENTS

We are thankful to Zhibin Lu and Carl Virtanen for helping us set up CReSCENT’s environment at the HPC4Health system. This project is supported in part by the Canadian Center for Computational Genomics (C3G). C3G is part of the Genome Technology Platform (GTP), funded by Genome Canada through Genome Quebec and Ontario Genomics. TJP holds the Canada Research Chair in Translational Genomics and is supported by a Senior Investigator Award from the Ontario Institute for Cancer Research and the Princess Margaret Cancer Foundation Gattuso-Slaight Personalized Cancer Medicine Fund.

